## Supplementary file 3 for "Evolutionarily conserved non-protein-coding regions in the chicken genome harbor functionally important variation"

### Supplementary Figures

**Figure S1. Distribution of conserved elements (CEs) along the chicken genome.** The barplot displays the fraction of the genome per chromosome covered by conserved elements.

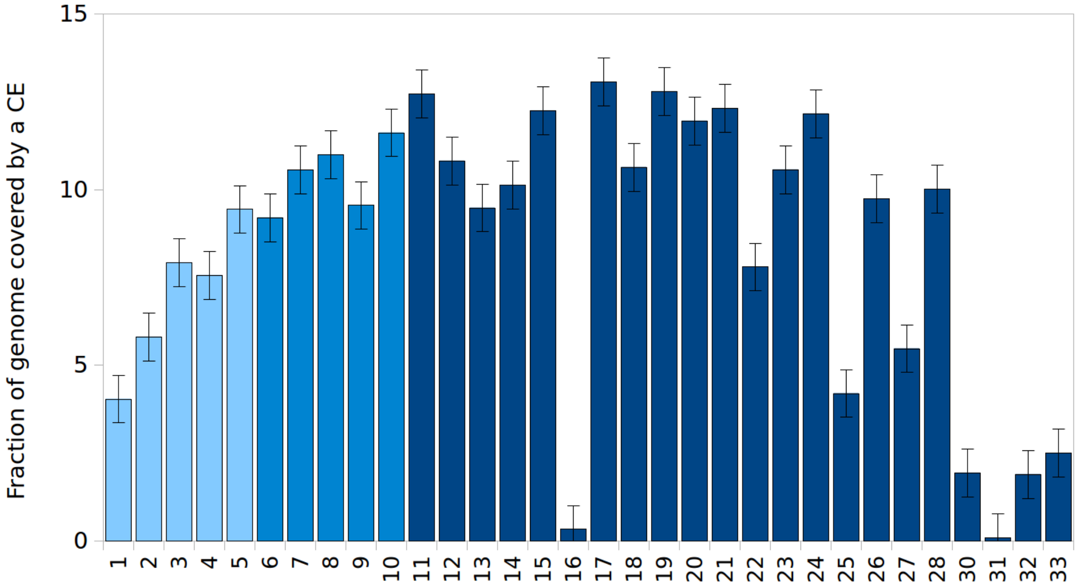

**Figure S2. Frequency size distribution of predicted conserved elements.** The y-axis shows the frequency, while the x-axis the size in base pairs (bp) of the predicted conserved elements.

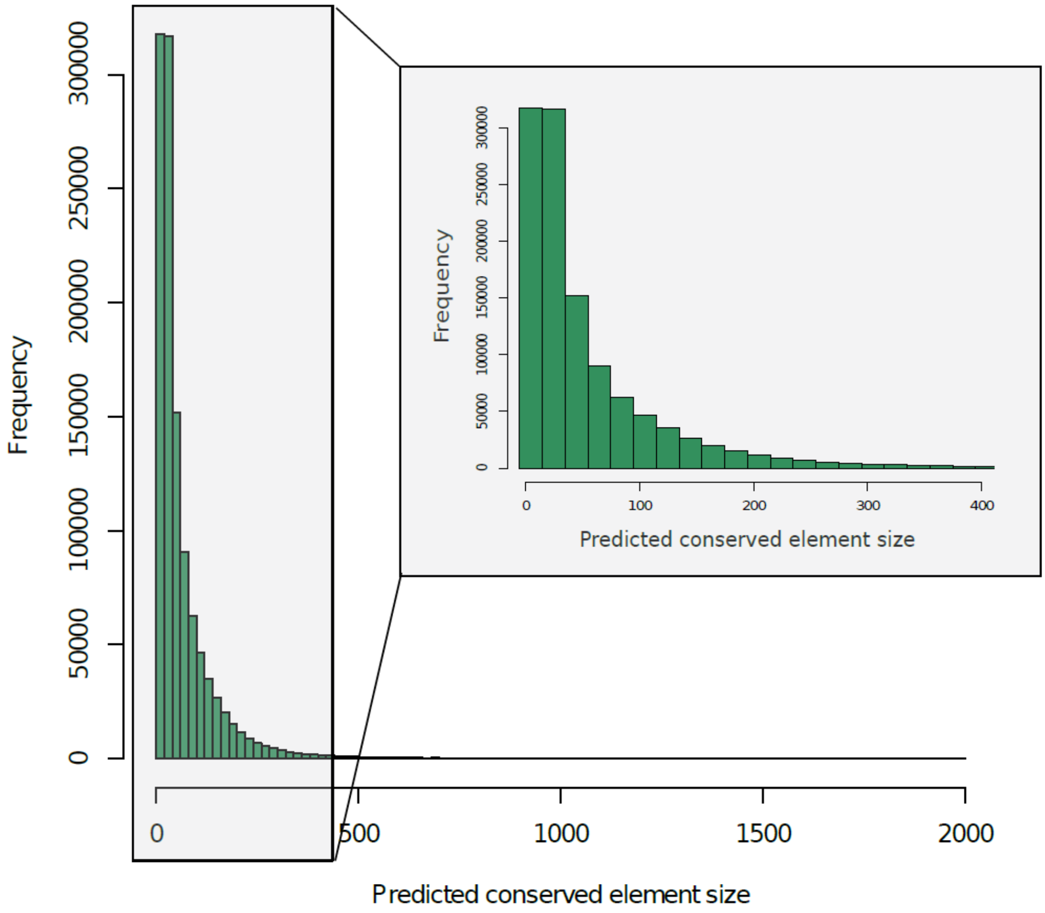

**Figure S3. Frequency size distribution of predicted conserved elements overlapping exonic-associated gene annotations.** The exonic-associated conserved elements include CDS, 5’UTR, 3’UTR, and promoter regions.

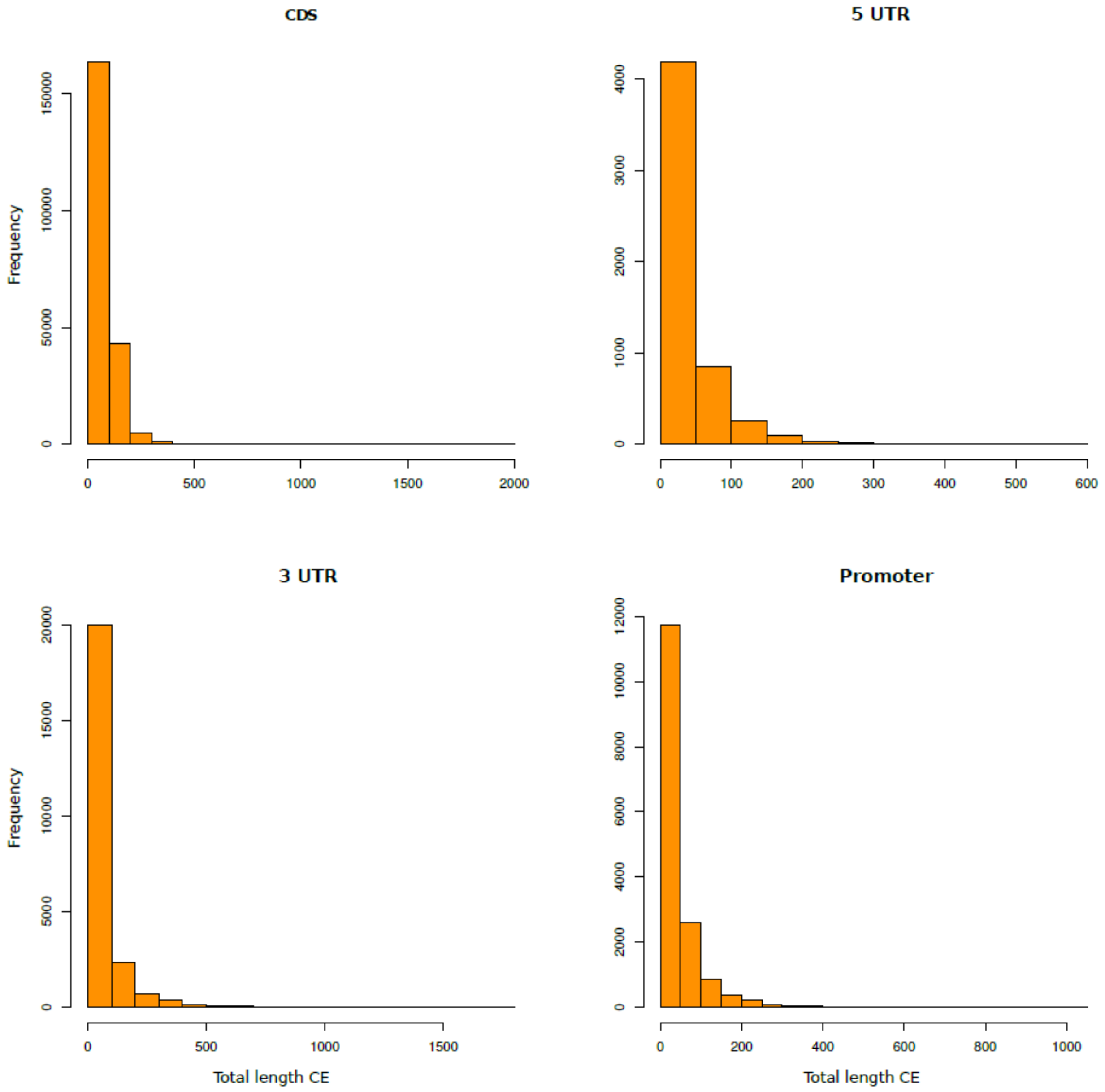

**Figure S4. Frequency size distribution of predicted conserved elements overlapping non-protein-coding gene annotations**. The non-protein-coding gene annotations include introns, lncRNA, and intergenic regions.

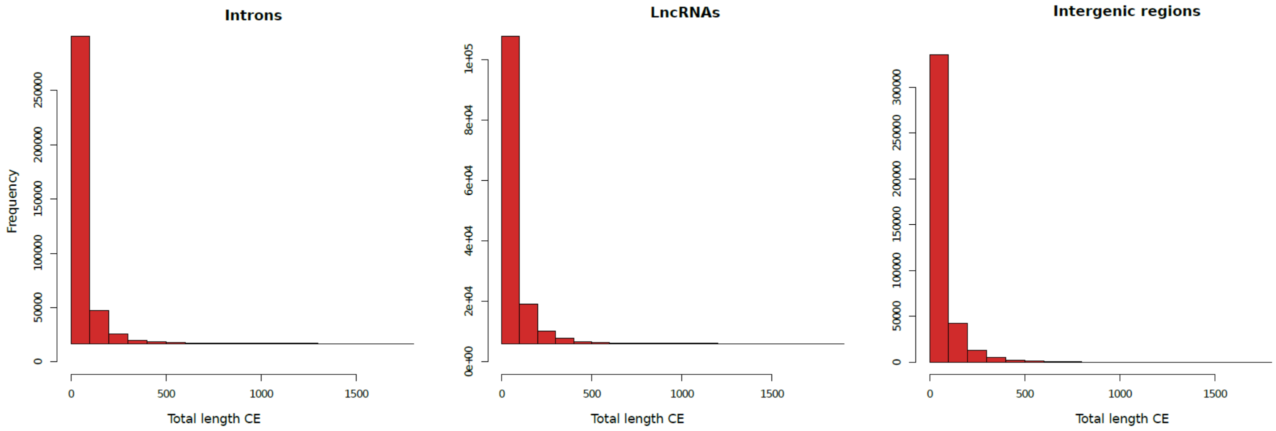

**Figure S5. Model performances measured in Receiver Operator Area under the Curve (ROC-AUC) and log-loss for three different ridge penalization terms (0.1, 1.0, 10.0)**. The scale is adjusted to make the differences between the models visible. Penalization of 1 was selected due to the lowest log-loss and largest ROC-AUC.

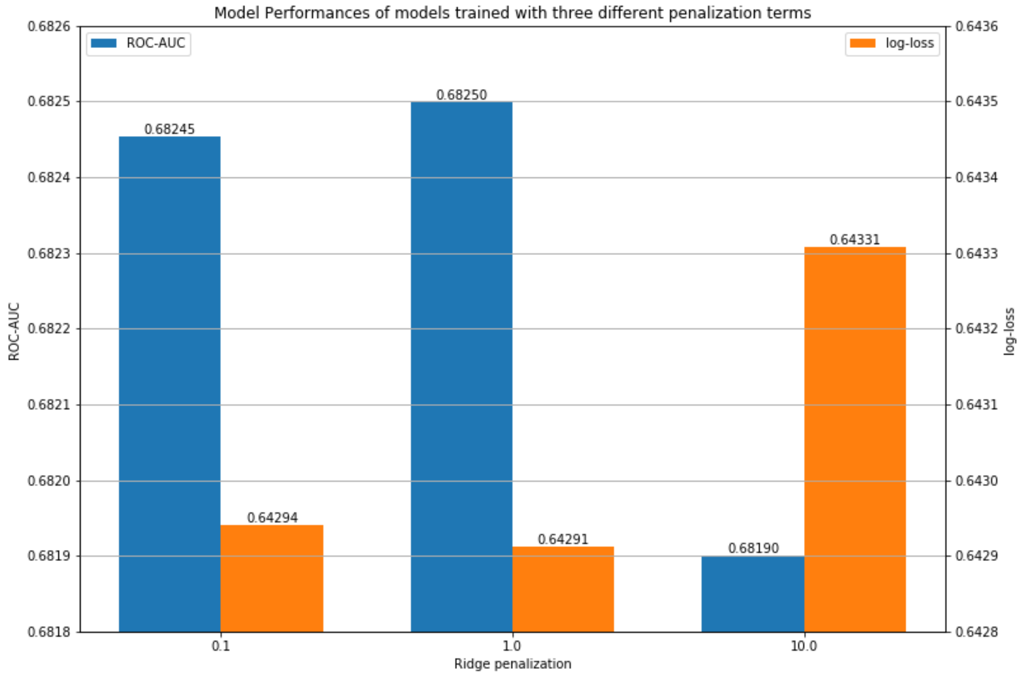

**Figure S6. chCADD score distribution of SNPs per VEP cateogy.** SNPs from the 169 chickens are categorized based on the VEP categories reported in Table S1 (SG: Stop-gained; CS: Canonical Splice; NS: Non-Synonymous; SN: Synonymous; SL: STOP-Lost; S: Splice Site; U5: 5’-UTR; U3: 3’-UTR; IG: Intergenic; NC: Noncoding-change; I: Intronic; UP: Upstream; DN: Downstream; O: Other). The label indicates the category and the number of SNPs falling into that category.

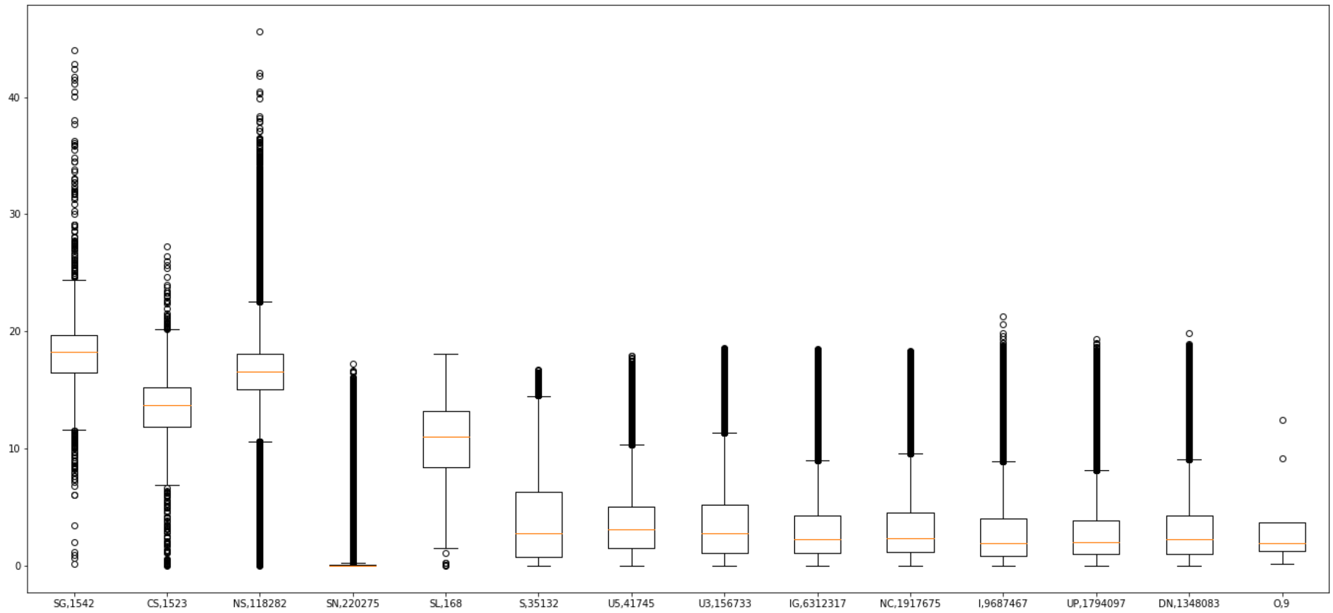

### Supplementary Table

**Table S1. VEP consequences summarized in 14 categories.** If multiple annotations exist for the same variant, the consequence is selected according to the displayed hierarchy, starting at 1 and ending at 14.

| Hierarchy | Abbreviation | VEP Consequence |
|---|---|---|
| 1 | SG | Stop Gained |
| 2 | CS | Canonical Splice |
| 3 | NS | Non-Synonymous |
| 4 | SN | Synonymous |
| 5 | SL | STOP Lost |
| 6 | S | Splice Site |
| 7 | U5 | 5’-UTR |
| 8 | U3 | 3’-UTR |

| | | |
|---|---|---|
| 9 | IG | Intergenic |
| 10 | NC | Noncoding-change |
| 11 | I | Intronic |
| 12 | UP | Upstream |
| 13 | DN | Downstream |
| 14 | O | Unknown / Other |

**Table S2. Top 10 Model features with the largest assigned weight and their explanations**

| Label | Model weight assigned to feature | Feature explanation |
|---|---|---|
| GerpS | 0.152568 | GERP rejected substitution score |
| 4PhCons_noChick | 0.128726 | 4-sauropsids PhastCons scores (excluding chicken) |
| I_GerpS | 0.109099 | GERP rejected substitution score for intronic sites |
| I_4PhCons_noChick | 0.0899441 | 4-sauropsids PhastCons scores (excluding chicken) for intronic sites |
| dnaProT | 0.083813 | DNA secondary structure prediction for ProT |
| 77PhCons_noChick | 0.0790709 | 4-amniota PhastCons scores (excluding chicken |
| dnaRoll | 0.0733429 | DNA secondary structure prediction for Roll |
| IG_4PhCons_noChick | 0.067539 | 4-sauropsids PhastCons scores (excluding chicken) for intergenic sites |
| I_dnaProT | 0.0671401 | DNA secondary structure prediction for ProT for intronic sites |
| IG_GerpS | 0.0635293 | GERP rejected substitution score for intergenic sites |

**Table S3. List of annotations which form the set of descriptive features for which model weights are learned.** Missing values are imputed via the specified values. Annotations of the type (factor) are OneHotEncoded and combinations between annotations form the final feature set.

| Annotation label | Data type | Imputed value | Annotation description |
|---|---|---|---|
| Ref | factor | | Reference allele |
| Alt | factor | | Observed allele |
| isTv | bool | 0.5 | Is transversion? |
| Consequence | factor | | VEP Consequence summaries |
| GC | num | 0.4 | Percent GC in a window of +/- 75bp |

| | | | |
|---|---|---|---|
| CpG | num | 0.02 | Percent CpG in a window of +/- 75bp |
| motifECount | int | 0.0 | Total number of overlapping motifs |
| motifEHIPos | bool | False | Is the position considered highly informative for an overlapping motif by VEP |
| motifEScoreChng | num | 0.0 | VEP score change for the overlapping motif site |
| Domain | factor | UD | Domain annotation inferred from VEP annotation (ncoils, tmhmm, sigp, lcompl, ndomain = "other named domain") |
| Dst2Splice | int | 0.0 | Distance to splice site in 20bp; positive: exonic, negative: intronic |
| Dst2SplType | factor | UD | Closest splice site is ACCEPTOR or DONOR |
| oAA | factor | UD | Amino acid of observed variant |
| nAA | factor | UD | Reference amino acid |
| Grantham | int | 0.0 | Grantham score: oAA,nAA |
| SIFTcat | factor | UD | SIFT category of change |
| SIFTval | num | 0.0 | SIFT score |
| cDNApos | int | 0.0 | Base position from transcription start |
| relcDNApos | num | 0.0 | Relative position in transcript |
| CDSpos | int | 0.0 | Base position from coding start |
| relCDSpos | num | 0.0 | Relative position in coding sequence |
| protPos | int | 0.0 | Amino acid position from coding start |
| relProtPos | num | 0.0 | Relative position in protein codon |
| dnaRoll | num | 0.23 | Predicted local DNA structure effect on dnaRoll |
| dnaProT | num | 0.68 | Predicted local DNA structure effect on dnaProT |
| dnaMGW | num | 0.03 | Predicted local DNA structure effect on dnaMGW |
| dnaHelT | num | -0.12 | Predicted local DNA structure effect on dnaHelT |
| GerpS | num | -0.17 | Rejected Substitution' score defined by GERP++ |
| GerpN | num | 0.64 | Neutral evolution score defined by GERP++ |
| GerpRS | num | 0.0 | Gerp element score |
| GerpRSpval | num | 1.0 | Gerp element p-Value |
| 4PhCons_noChick | num | 0.17 | 4-taxa-sauropsids PhastCons score (excl. chicken) |
| 37PhCons_noChick | num | 0.13 | 37-taxa-Amniota PhastCons score (excl. chicken) |
| 77PhCons_noChick | num | 0.2 | 77-taxa-Vertebrate PhastCons score (excl. chicken) |
| 4PhyloP_noChick | num | 0.07 | 4-taxa-sauropsids PhyloP score (excl. chicken) |
| 37PhyloP_noChick | num | 0.04 | 37-taxa-Amniota PhyloP score (excl. chicken) |
| 77PhyloP_noChick | num | 0.25 | 77-taxa-Vertebrate PhyloP score (excl. chicken) |
| minDistTSS | int | 10000000 | Distance to closest Transcribed Sequence Start (TSS) |

| | | | |
|---|---|---|---|
| minDistTSE | int | 10000000 | Distance to closest Transcribed Sequence End (TSE) |
| interaction-score | num | 0 | Interaction score from Hi-C interaction maps |
| Exp-score | int | 0 | RNA expression scores |
| Exp-pval | num | 1 | p-Value of RNA expression scores |
| Exp-logFC | num | 0 | Log-Fold change of RNA expression |
| OChrom-Peaknb | Int | 0 | Read number for open Chromatin; ATAC-seq |
| OChrom-pval | num | 1 | p-Value for open chromatin; ATAC-seq |
| OChrom-logFC | num | 0 | Log-Fold change for ATAC-seq |

**Table S4. Phenotypes of homologous genes of the top 10 intronic CNEs.** The top 10 intronic CNEs were selected based on the largest differences between the 1$^{st}$ and 3$^{rd}$ to the 2$^{nd}$ section within a CNE.

| **Chr** | **Start CE** | **End CE** | **Ensembl ID** | **Human phenotype** | **Mouse phenotype** | **Rat phenotype** |
|---|---|---|---|---|---|---|
| 10 | 3714893 | 3714924 | ENSGALG00000002883 | Autosomal Recessive Mental Retardation, intellectual developmental disorder and retinitis pigmentosa, Retinitis pigmentosa | abnormal heart left ventricle morphology, decreased grip strength, decreased large unstained cell number, decreased lean body mass, increased or absent threshold for auditory brainstem response, male infertility, preweaning lethality incomplete penetrance | - |

| | | | | | | |
|---|---|---|---|---|---|---|
| 3 | 21413813 | 21413883 | ENSGALG00000009791 | - | abnormal endocrine pancreas morphology, abnormal eye development, abnormal lens development, abnormal lens morphology, abnormal liver development, abnormal lymph organ development, absent horizontal cells, decreased hepatocyte proliferation, decreased lymphatic vessel endothelial cell number, edema, increased pancreatic acinar cell number, lethality throughout fetal growth and development complete penetrance, no abnormal phenotype detected, small liver, small pancreas | Status Epilepticus |
| 5 | 46655335 | 46655367 | ENSGALG00000011093 | Non-syndromic male infertility due to sperm motility disorder, spermatogenic failure 27 | abnormal cerebellum morphology, abnormal head shape, abnormal internal nares morphology, abnormal respiratory motile cilium morphology, abnormal respiratory motile cilium physiology, abnormal sperm head morphology, arrest of spermatogenesi | - |

| | | | | | | |
|---|---|---|---|---|---|---|
| | | | | | s, azoospermia, enlarged lateral ventricles, hydroencephaly, impaired mucociliary clearance, male infertility, oligozoospermia, postnatal growth retardation, premature death, respiratory system inflammation, rhinitis, thin cerebral cortex | |
| 5 | 29339796 | 29339843 | ENSGALG00000009587 | Hereditary hyperekplexia, hyperekplexia 1, Molybdenum cofactor deficiency complementation group C, Sulfite oxidase deficiency due to molybdenum cofactor deficiency type C | abnormal axon extension, abnormal motor neuron morphology, abnormal nervous system electrophysiology, abnormal neuromuscular synapse morphology, abnormal posture, abnormal retinal inner plexiform layer morphology, abnormal suckling behavior, abnormal vocalization, apnea, decreased motor neuron number, hyperresponsive, increased motor neuron number, motor neuron degeneration, neonatal lethality complete penetrance, no abnormal phenotype detected | inherited metabolic disorder |

| | | | | | | |
|---|---|---|---|---|---|---|
| 1 | 66913273 | 66913298 | ENSGALG00000013244 | Acromegaloid facial appearance syndrome, atrial fibrillation familial 12, brugada syndrome, cantu syndrome, cantu syndrome hypertrichotic osteochondrodysplasia, cardiomyopathy dilated 1o, familial atrial fibrillation, Familial isolated dilated cardiomyopathy, Hypertrichosis-acromegaloid facial appearance syndrome, Hypertrichotic osteochondrodysplasia Cantu type | abnormal ST segment, abnormal systemic arterial blood pressure, abnormal vascular smooth muscle physiology, abnormal vasoconstriction, artery stenosis, hypertension, hypoglycemia, improved glucose tolerance, increased insulin sensitivity, increased muscle cell glucose uptake, increased systemic arterial diastolic blood pressure, increased systemic arterial systolic blood pressure, premature death, slow postnatal weight gain | Diabetes Mellitus Experimental, hypertension, Parkinsonian Disorders, Sciatic Neuropathy, Ventricular Fibrillation, Ventricular Tachycardia |
| 18 | 2616518 | 2616533 | ENSGALG00000001375 | isolated cytochrome c oxidase deficiency, leigh syndrome, mitochondrial complex iv deficiency | - | - |
| 3 | 46314957 | 46314976 | ENSGALG00000012256 | - | abnormal miniature endplate potential, abnormal nervous system physiology, abnormal neuromuscular synapse morphology, increased sensitivity to xenobiotic induced morbidity/mortality | Duchenne muscular dystrophy, Ovarian Neoplasms |

| | | | | | | |
|---|---|---|---|---|---|---|
| 1 | 34590834 | 34590912 | ENSGALG00000009895 | Fraser syndrome, Fraser syndrome 3 | abnormal blood coagulation, abnormal blood vessel morphology, abnormal brain morphology, abnormal corneal epithelium morphology, abnormal corneal stroma morphology, abnormal cornea thickness, abnormal eye development, abnormal eyelid morphology, abnormal eye morphology, abnormal iris morphology, abnormal kidney development, abnormal lens development, abnormal lens vesicle development, abnormal limb morphology, abnormal neural tube morphology, abnormal retina morphology, absent kidney, absent limbs, anophthalmia, aphakia, bleb, blistering, cataract, clubfoot, corneal opacity, decreased body size, embryonic lethality during organogenesis incomplete penetrance, eye hemorrhage, eyelids open at birth, hemorrhage, interdigital webbing, intracranial | - |

| | | | | | | |
|---|---|---|---|---|---|---|
| | | | | | hemorrhage, kidney cysts, microphthalmia, open neural tube, perinatal lethality incomplete penetrance, polycystic kidney, polydactyly, prenatal lethality complete penetrance, single kidney, small kidney, syndactyly | |
| 2 | 102926228 | 102926256 | ENSGALG00000014998 | - | abnormal CNS glial cell morphology, abnormal endometrial gland morphology, abnormal endometrium morphology, absent corpus callosum, decreased litter size, dilated uterus, endometrium hyperplasia, enlarged uterus, increased endometrial carcinoma incidence, reduced female fertility | - |

| | | | | | |
|---|---|---|---|---|---|
| 7 | 26190151 | 26190213 | ENSGALG00000011645 | | preweaning lethality incomplete penetrance |
